# Clonal expansion of mitochondrial DNA deletions is a private mechanism of ageing in long-lived animals

**DOI:** 10.1101/273326

**Authors:** Lakshmi Narayanan Lakshmanan, Zhuangli Yee, Li Fang Ng, Rudiyanto Gunawan, Barry Halliwell, Jan Gruber

## Abstract

Disruption of mitochondrial metabolism and loss of mitochondrial DNA (mtDNA) integrity are widely considered as evolutionarily conserved (public) mechanisms of ageing (López-Otín *et al.* 2013). Human ageing is associated with loss in skeletal muscle mass and function (Sarcopenia), contributing significantly to morbidity and mortality. Muscle ageing is associated with loss of mtDNA integrity. In humans, clonally expanded mtDNA deletions co-localize with sites of fiber-breakage and atrophy in skeletal muscle. mtDNA deletions may therefore play an important, possibly causal role in sarcopenia. The nematode *Caenorhabditis elegans* also exhibits age-dependent decline in mitochondrial function and a form of sarcopenia. However, it is unclear if mtDNA deletions play a role in *C. elegans* ageing. Here we report identification of 266 novel mtDNA deletions in ageing nematodes. Analysis of the mtDNA mutation spectrum and quantification of mutation burden indicates that (1) mtDNA deletions in nematode is extremely rare, (2) there is no significant age-dependent increase in mtDNA deletions and (3) there is little evidence for clonal expansion driving mtDNA deletion dynamics. Thus, mtDNA deletions are unlikely to drive the age-dependent functional decline commonly observed in *C. elegans*. Computational modelling of mtDNA dynamics in *C. elegans* indicates that the lifespan of short-lived animals such as *C. elegans* is likely too short to allow for significant clonal expansion of mtDNA deletions. Together, these findings suggest that clonal expansion of mtDNA deletions is likely a private mechanism of ageing predominantly relevant in long-lived animals such as humans and rhesus monkey and possibly in rodents.

## Introduction

Eukaryotic cells typically contain connected networks of mitochondria, each containing multiple redundant copies of mitochondrial DNA (mtDNA). The copy-number of mtDNA per cell is highly dependent on cell type and energy requirements but usually cells contain 1000-10000s of mtDNA (Miller *et al.* 2003). The nematode *Caenorhabditis elegans (C. elegans)*, a widely used model organism in developmental biology, neurobiology and ageing, comprises 959 cells, together typically containing between 200,000 and 300,000 mtDNA molecules (Tsang & Lemire 2002a; Gruber *et al.* 2011). The copy number of mtDNA per animal declines with age, with aged nematodes containing almost 30% fewer mtDNA compared to young adults (Gruber *et al.* 2011). Errors during mtDNA replication or during repair of DNA damage can result in mtDNA mutations, including large deletions spanning multiple mtDNA genes (Krishnan *et al.* 2008; Alexeyev *et al.* 2013). Mutant mtDNA molecules can be amplified during normal mtDNA maintenance, creating additional copies of the mutant sequence within the same cell (Stewart & Chinnery 2015). During clonal expansion, mutant mtDNA increases in abundance relative to the wildtype (WT) sequence (Stewart & Chinnery 2015). When the fraction of mutant mtDNA exceeds ~60%, the excess mutant mtDNA can cause mitochondrial dysfunction, cell atrophy or death (Wanagat *et al.* 2001; Rossignol *et al.* 2003).

Highly clonally expanded mtDNA deletions were first detected in muscle fibers of human myopathy patients (Holt *et al.* 1988). Later studies confirmed that clonal mtDNA deletions are also present in muscle of healthy, aged humans, where they co-localize with zones of atrophy of muscle fibers and fiber breakage (Bua *et al.* 2006). In addition to muscle, amplified mtDNA deletions have been reported in substantia nigra neurons in Parkinson disease patients, suggesting their involvement in some aspects of ageing and disease processes in brain as well (Bender *et al.* 2006). While clonally expanded mtDNA mutations are detected most frequently in post mitotic tissues such as brain and muscle, they may also occur in mitotic tissues such as in colonic epithelium and crypt stem cells (McDonald *et al.* 2006; Baines *et al.* 2014). These observations support a role of mtDNA mutations in age-associated functional decline and disease, at least in humans.

Model organisms, especially mice, rats, flies, nematodes and yeast, are frequently utilized to accelerate discovery of mechanisms of ageing and to identify targets for intervention. However, data on mtDNA deletion in simple model organisms are relatively sparse and it is unclear if mtDNA deletion accumulation is an evolutionarily conserved, “public” mechanism of ageing (Partridge & Gems 2002; Gruber *et al.* 2015). On the one hand, mtDNA deletions have been reported in tissues from rhesus monkeys (Gokey *et al.* 2004), mice (Chung *et al.* 1996), rats (Pak *et al.* 2005), flies (Yui & Matsuura 2006) and nematodes (Melov *et al.* 1994; Melov *et al.* 1995). On the other hand, at least two studies have questioned whether mtDNA deletions are commonly present in aged mice, finding levels in aged mice undetectably low (Williams *et al.* 2010; Ameur *et al.* 2011). Little is known about the role of mtDNA deletion accumulation in nematode ageing. Similar to other animals, nematodes experience significant hypo-metabolism and decline in mitochondrial function as they age. Oxygen consumption and ATP levels (indicators of metabolism) significantly decline with age in nematodes (Braeckman *et al.* 2002; Yasuda *et al.* 2006; Gruber *et al.* 2011). Behaviors based on muscular and nervous system function, including pharyngeal pumping, spontaneous movement and motility type, also decline rapidly with age in *C. elegans* (Herndon et al. 2002; Fisher 2004; Collins et al. 2008; Gruber et al. 2011). A significant decline in muscle function and structural integrity is a prominent feature of ageing in *C. elegans* (Herndon et al. 2002; Fisher 2004; Collins et al. 2008; Gruber et al. 2011). This age-dependent deterioration of the *C. elegans* musculature resembles sarcopenia, involving muscle cell shrinkage, fraying and loss of structure of muscle fibers with loosening of packing geometry of sarcomere fibers (Herndon *et al.* 2002). Ageing in *C. elegans* is also associated with increased oxidative damage and loss of structural integrity of muscle mitochondria (Yasuda *et al.* 2006).

However, there are only two studies that directly report an age-associated increase in mtDNA deletions abundance in *C. elegans* (Melov *et al.* 1994; Melov *et al.* 1995). Using single primer pairs to detect mtDNA deletions, Melov *et al.* were able to show that mtDNA deletions are more readily detectable in old than in young nematodes (Melov *et al.* 1994; Melov *et al.* 1995). However, no absolute or relative quantification mtDNA copy number can be derived from these data and, all in all, only 6 age-associated mtDNA deletions in *C. elegans* have previously been sequenced, meaning that no representative mutation spectrum can be derived either. Practically no information exists regarding the functional relevance (if any) of such deletions. It is, therefore, unclear if mtDNA deletions ever become sufficiently abundant to be a plausible cause of the hypo-metabolism and sarcopenia phenotypes of ageing nematode. Indeed, there are reasons to question if clonal expansion should be expected to occur commonly in short-lived animals such as *C. elegans*. In humans, mtDNA deletions are detected late in life, typically at > 50 years of age, by which age, atrophied muscle fibers are found to be enriched with clonally expanded mtDNA deletions (Bua *et al.* 2006). By contrast, nematodes can exhibit signs of sarcopenia as early as 7 days after hatching (Herndon *et al.* 2002; Fisher 2004). We have previously shown that mtDNA maintenance, in particular mtDNA turnover, is a crucial determinant of the dynamics of mutant mtDNA accumulation (Poovathingal *et al.* 2009; Poovathingal *et al.* 2012; Tam *et al.* 2013; Tam *et al.* 2015). Given that clonal expansion takes decades to occur in human muscle, we questioned if these processes scale to allow clonal expansion to occur over only seven days in *C. elegans*. Here, we address the following questions: (1) what is the typical mtDNA deletion mutation burden in young and old worms? (2) Do mtDNA deletions in nematodes accumulate to physiologically significant levels over the course of life? (3) Are mtDNA deletions a plausible cause of age-dependent decline in metabolism and sarcopenia in *C. elegans*? (4) Is there evidence for significant clonal expansion of mtDNA deletions in *C. elegans*? (5) What sequence features are associated with mtDNA deletions in *C. elegans* compared to mammals? And finally, (6) from a dynamical perspective, should we expect clonal expansion of mtDNA deletions within the short lifespan of worms and, by extension, of other short-lived model organisms? To address these questions, we use two complementary approaches. Experimentally, we detect (capture), sequence and quantify mtDNA deletion mutations in cohorts of ageing nematodes. Using this approach, we find that mtDNA deletions remain extremely low in abundance even in old animals. Based on comparing samples within and between biological replicates, we find little evidence for the hypothesis that the detected mtDNA mutations result from significant clonal expansion. Finally, using our previously developed stochastic mathematical model of mtDNA deletions dynamics, we show that clonal expansion is unlikely to occur commonly in *C. elegans* given its short lifespan.

## Results

### Survival analysis of nematode cohorts

Our aim was to evaluate age-dependent changes in mtDNA integrity in *C. elegans* as a possible explanation of age-dependent functional and metabolic decline. A confounding factor in studies of mtDNA integrity in *C. elegans* is the presence of large amounts of mtDNA in oocytes, eggs and juvenile offspring during the reproductive phase of life (Tsang & Lemire 2002a). To avoid this confounder, we used nematodes of a germline proliferation deficient (*glp-1*) strain for all of our experiments. Nematodes of this strain are sterile at the restrictive temperature of 25°C and do not produce or lay eggs at this temperature. Mutations in *glp-1* are genetic equivalents of germline ablation. At the restrictive temperature of 25°C, the *glp-1* strain is sterile and (relative to WT at 25°C) long-lived but its somatic development has been reported to be normal (Tsang & Lemire 2003). Also, direct comparison of metabolic ageing between *glp-1* and WT (N2) shows that the trajectories of *glp-1* and WT are similar (Shoyama *et al.* 2009).

**Figure 1.**
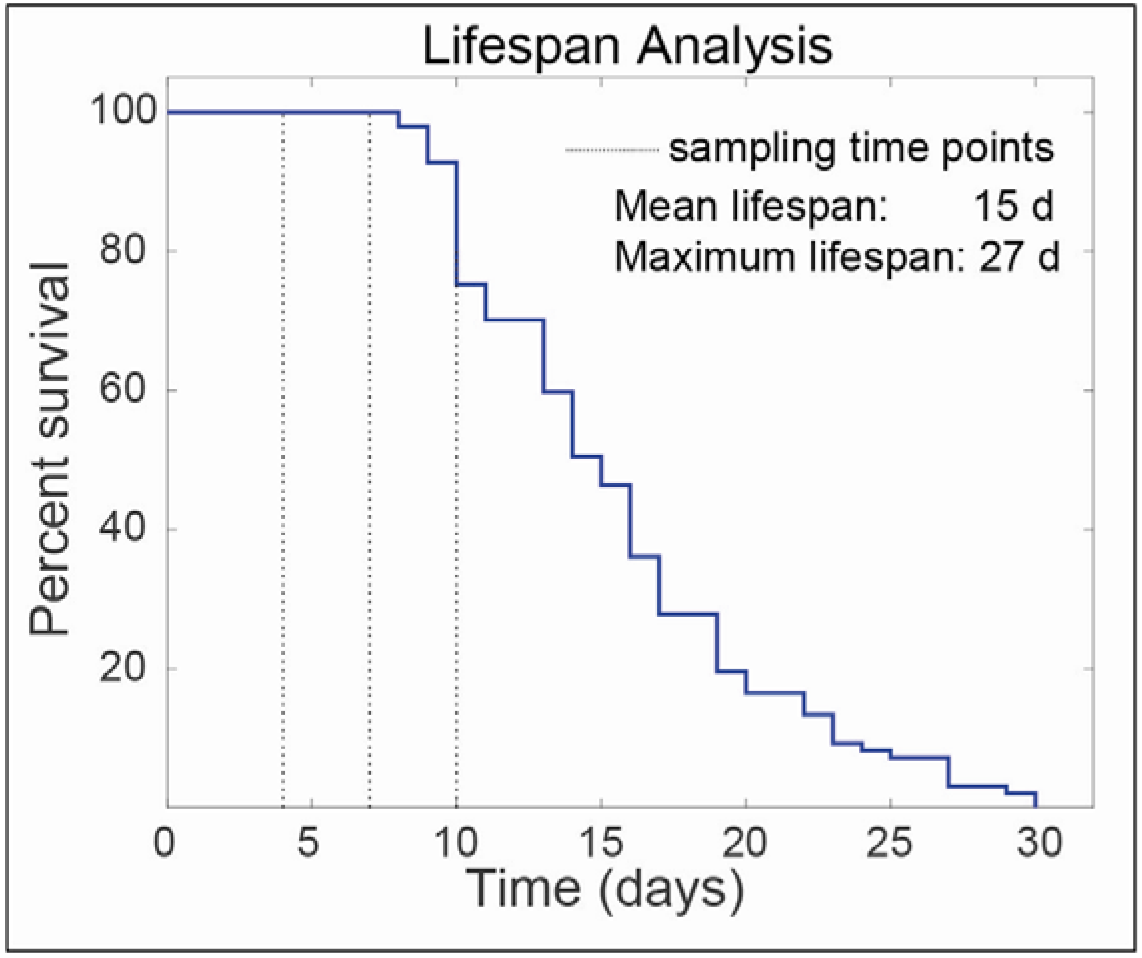
Typical survival plot for *C. elegans glp-1* strain at 25°C in our laboratory. Data shows survival data for *n* = 97 nematodes from a cohort maintained at 25°C. Significant increases in morbidity and cumulative mortality were observed after day 10. To avoid confounders such as dead worms and selection effects during the mutation capture experiments, cohorts were sampled on days 4 (3 cohorts), 7 (5 cohorts) and 10 (3 cohorts) for mtDNA extraction.

In order to determine the appropriate time points to sample mtDNA from the cohort we determined the survival of a cohort of *glp-1* mutants in our lab at the restrictive temperature of 25 °C. Median lifespan was 15 days and the maximum lifespan, defined as the average of the longest surviving 10% of the population, was 27 days (Fig. 1). Development from egg to young adult takes approximately 3 days and by day 4 of life, all animals are adults. While *glp-1* worms are sterile at the restrictive temperature, peak reproductive and physical fitness in *C. elegans* is typically reached between day 6 and 8. In our cohort, there was a significant increase in cumulative mortality after day 10 of life with few worms dying before this age (Fig. 1), meaning that day 10 was the last day on which large numbers of living animals for analysis could be readily obtained. Furthermore, we and other have evaluated mitochondrial, metabolic and mitochondrial ageing trajectories in *glp-1*. At 25°C, oxygen consumption by *glp-1* declines significantly from a peak between day 3 and 4 and is substantially diminished (to less than 30% of the peak value) by day 10 to 12 (Shoyama *et al.* 2009; Gruber *et al.* 2011). This dramatic decline in oxygen consumption is associated with reduced energy levels, as evaluated by steady state ATP concentration, as well as loss of mitochondrial content and increased oxidative damage to protein and mtDNA (Gruber *et al.* 2011). By day 10, *glp-1* at 25°C exhibit clear evidence of mitochondrial decline (Shoyama *et al.* 2009; Gruber *et al.* 2011). We therefore chose three sampling time points, days 4, 7 and 10 to represent young healthy worms, worms at around the peak reproductive age and aged worms subject to significant functional decline and experiencing increasing mortality.

**Figure 2.**
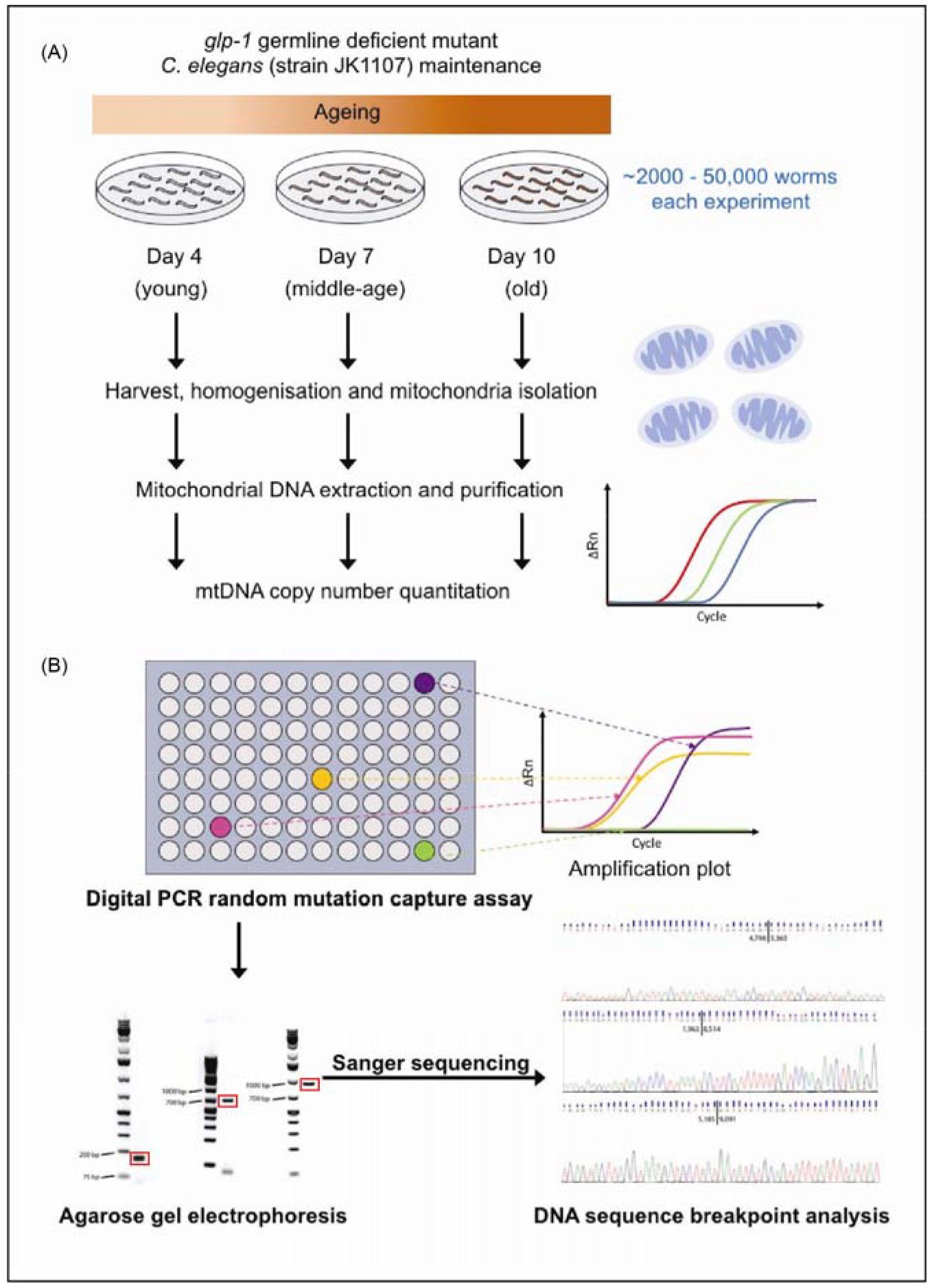
Schematic overview of the mutation capture assay. (A) Nematodes were sampled from ageing cohorts at ages 4, 7 and 10 days. Differential centrifugation and DNA extraction were used to obtain mtDNA samples for each age group. mtDNA copy number in samples was then quantified using digital PCR. (B) For each age group, the presence of mtDNA deletions was detected using 236 distinct pairs of primers decorating ~65% of the total mtDNA sequence of *C. elegans*. mtDNA from PCR wells with positive amplification signals (sequence specific fluorescence probe) were further analyzed using gel electrophoresis and DNA sequencing to confirm the presence of mtDNA deletions and to identify breakpoints.

**Table 1.** Experimental design for mutation capture assay. Experimental design for mutation capture assay. We used at least three independent cohorts for each age group. Dependent on DNA yield, we then carried out between 1 and 4 repeat mutation capture assays per ageing cohort. For one of the day-10 cohorts we instead carried out a total of ten repeats of the mutation capture assay using a single tube of mtDNA extract from this single day-10 cohort. This was done to permit comparison of deletions detected repeatedly between cohorts and within a single cohort.

| Age group | Number of cohorts | Total replicates (mutation capture assays) | Total deletions detected | Average deletions per replicate |
| --- | --- | --- | --- | --- |
| day-4 | 3 | 11 | 101 | 9.20 |
| day-7 | 5 | 5 | 55 | 11.00 |
| day-10 | 3 | 12 | 156 | 13.00 |

### Random mutation capture assay for detecting mtDNA deletions

For each age group, we used at least 3 independent worm cohorts and at least 5 replicate mutation capture experiments (see Table 1). Detailed information about the cohort and replicate counts for each age group can be found in supporting information (Table S1). We then used a digital PCR-based mutation capture assay (Vogelstein & Kinzler 1999) to detect deletions in mtDNA samples obtained in mitochondrial preparations from cohorts of ageing *C. elegans* (Fig. 2). Each mutation capture experiment involved scanning for mtDNA deletions using a set of overlapping primer pairs. Specifically, we designed twelve forward and thirty-six reverse primers, forming a total of 236 possible primer pairs (see Methods). The primers were designed to decorate a 9000 bp long region (positions 1800 - 10800) within the *C. elegans* mtDNA sequence (Fig. 3). Primer positions were chosen within this segment of the mtDNA to minimize primer amplification bias and to obtain a representative sample of the *in vivo* mtDNA deletion spectrum. Primer pairs and RT-PCR protocol were designed such that for each primer pair, the amplicon length in the absence of significant deletions was larger than what could be amplified under the PCR conditions. Using Monte Carlo approach, we sampled 100,000 randomly generated deletion breakpoint positions between positions 1800 - 10800 in *C. elegans* mtDNA and confirmed that for 95% of deletions of length ≥ 500 bp, at least one of the 236 primer combinations was able to amplify the deletion (see Fig. S1, Supporting information). We carried out the mutation capture assay using one primer pair per well using three 96 well PCR plates (236 primer pairs plus positive and negative controls) per capture experiment. Samples were diluted to a final concentration of 10,000 mtDNA molecules per μl. Subsequently, 1μl (~10,000 molecules) of mtDNA was added to each of the 236 wells of the mutation capture assay. Through serial-dilution, spike-recovery experiments using an artificial deletion sequence (see Fig. S2, Supporting information), we verified that the presence of excess WT mtDNA did not interfere with amplification of the expected mutant amplicon even at very low (stochastic) mutant copy numbers (see Fig. S3, Supporting information). In each mutation capture experiment, we therefore scanned for mtDNA mutation using all 236 primer pairs, with each distinct primer pair in a separate well (Fig. 2). Since WT mtDNA cannot be amplified by the primer pairs, any amplified well indicates the presence of at least one mtDNA molecule in that well carrying a deletion large enough to permit amplification. Each well that showed amplification was therefore considered a putative deletion detection event. Each such putative deletion was gel purified and confirmed by sequencing. For each age group, we thus obtained a list of deletions and the number of times that each of the deletion sequences was detected among the replicate measurements from the mutation capture experiments (Table S2, Supporting information). The higher the abundance of a specific deletion in the mtDNA extract of a worm cohort, the higher will be the chance that this mutation is repeatedly detected in the replicate measurements. The raw results from these replicated mutation capture experiments (list of mutant sequences and counts of how many times each mutation was seen) contain both qualitative (mutation spectrum) and quantitative (abundance) information.

**Figure 3.**
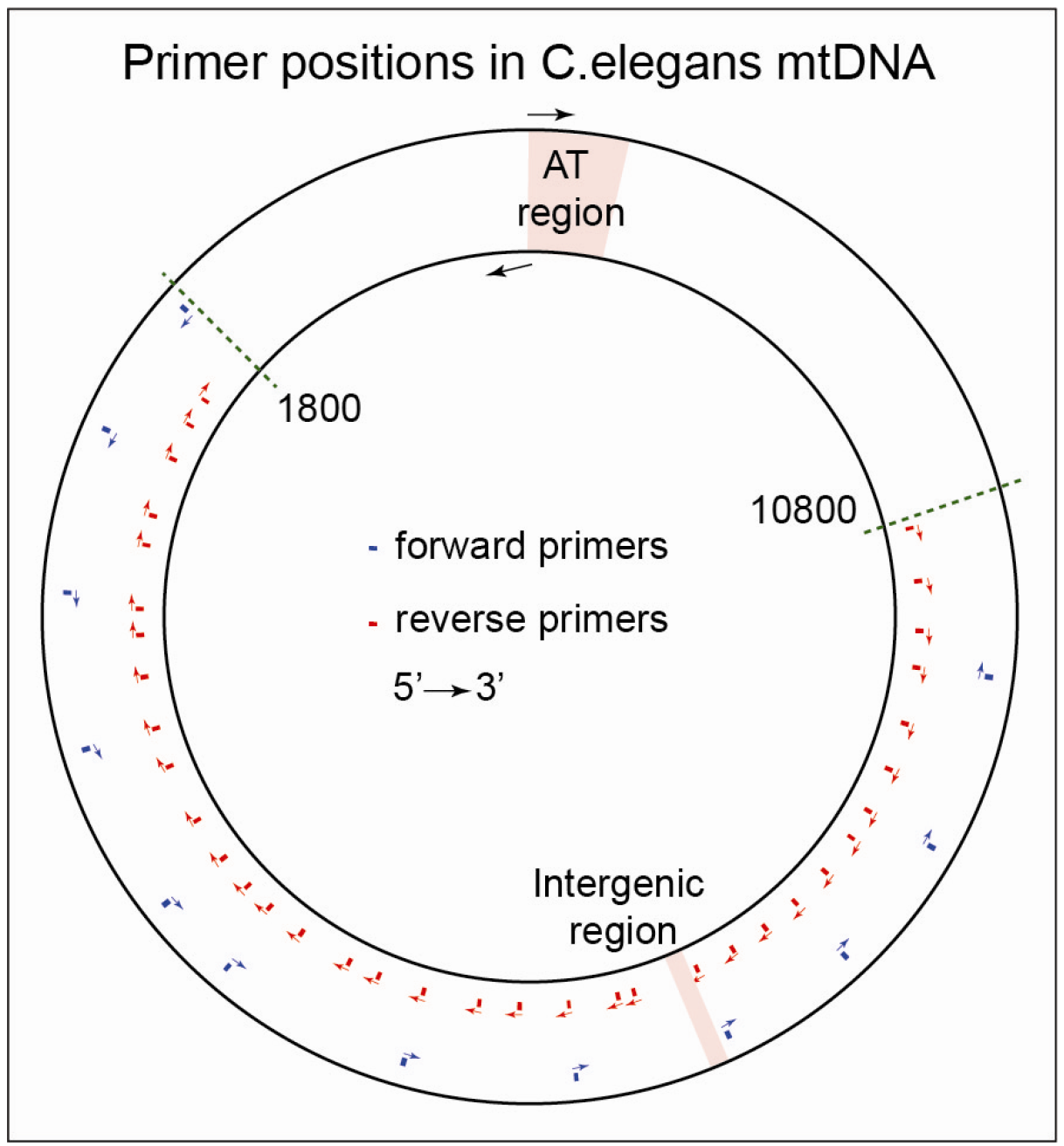
Locations of primers within the *C. elegans* mtDNA sequence. A 9000 bp region (positions 1800-10800) within *C. elegans* mtDNA was scanned for mtDNA deletions using 236 distinct forward-reverse primer pairs formed by 12 forward and 36 reverse primers. AT-rich region and intergenic region positions were obtained from mtDNA sequence annotation (NCBI accession number: X54252.1).

### Quantification of mutant burden of ageing associated mtDNA deletions

The data above are not easily comparable between experiments or with other reports in the literature without converting detection counts into mutant fraction values (i.e. ratio of mutant mtDNA count to total mtDNA count). Furthermore, mutation capture is a digital PCR technique and intrinsically stochastic in nature. To obtain quantitative information regarding age-dependent change in the abundance of mtDNA deletions it is necessary to relate the number of amplified wells from a given experiment with the mutation fraction (i.e. number of mutations per mtDNA molecule), while taking into account experimental design and sampling statistics. Due to the stochasticity associated with the sampling of mutant molecules during the different steps of the experiment, a same mutant fraction value in the starting sample can result in a distribution of well amplification counts. This effect is expected to be prominent when the copy number of mutants is low. Capturing the net effect of sampling statistics through serial dilution experiments using a known mutant molecule would require a large number of replicate experiments to capture the distribution of well amplification counts associated with each starting mutant fraction and hence would be prohibitive. For this reason, we developed a computational procedure to estimate the mutant fraction in mtDNA extracts based on the observed primer pair amplification data from the mutation capture assay. This method relies on Monte Carlo sampling methods (see Supporting information). Briefly, starting from an *in silico* worm homogenate with a specific mutant fraction, we simulated each step of the extraction procedure and mutation capture assay using a series of hypergeometric sampling events. Sample volumes and dilutions used for hypergeometric sampling were identical to the actual experimental values. For each hypothetical mutant fraction (virtual deletion burden), we repeated the entire *in silico* sampling experiment 1000 times. Using this procedure, we obtained an estimate of the distribution of observed amplification counts (the number of PCR wells amplified per experiment) that would be expected for each hypothetical mutant fraction.

By repeating this procedure for a wide range of plausible virtual mutant fractions, we generated a calibration curve relating the number of observed amplification events (wells containing at least one mtDNA deletion) during the mutation capture experiments with the homogenate mutant fraction (Fig. S4, Supporting information). Using this calibration curve to interpret the experimentally observed number of amplified wells from each mutant capture experiment, we estimated the most likely actual worm homogenate mutant fraction for each age group as well as a 95% confidence interval for these estimates. A detailed description of the computational method is provided in the supporting information. Using this approach, we quantified the mutant fraction in nematode cohorts at age 4, 7 and 10 days (Fig. 4). Analyzing the mutation fraction for these different cohorts we observed a slight trend towards higher mutation burden with age, but there is no statistically significant difference between young and old animals (Linear regression, zero value for slope cannot be rejected, *p*-value: 0.584).

**Figure 4.**
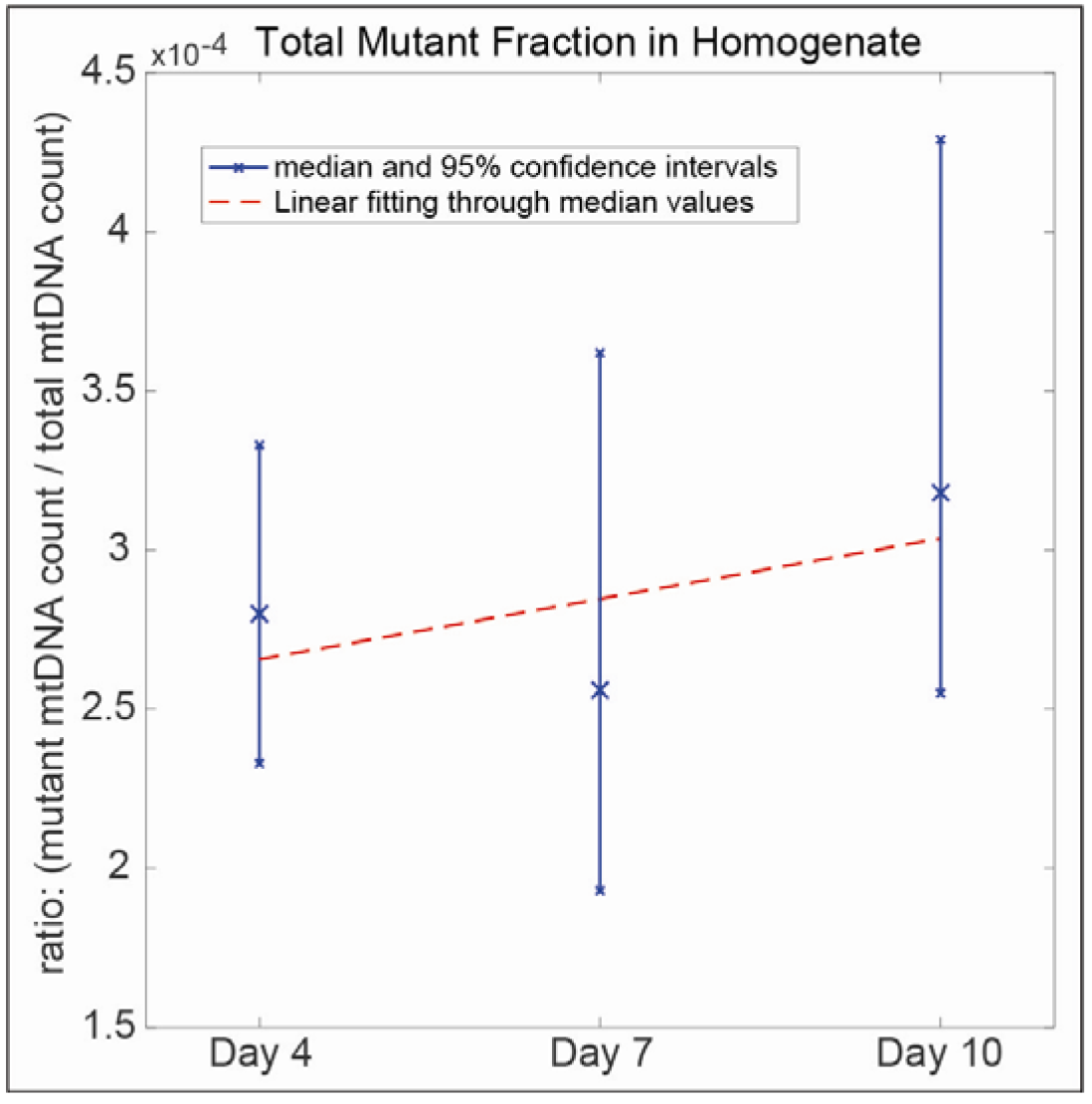
Experimentally determined mtDNA deletion abundance as mutant fraction relative to WT. Mutant fraction values were estimated for each age group from mutation capture assay results using Monte Carlo modelling of sampling statistics (see Fig. S2, Supporting information). Median mutant fraction and 95% confidence intervals (blue) in worm homogenates of cohorts at different ages are shown together with a linear regression best-fit line (red) of the same data. The trend towards higher mutant fraction in older cohorts is non-significant (a slope of zero cannot be rejected, p-value = 0.584).

To evaluate the physiological relevance (or lack thereof) of these mtDNA deletion burdens in ageing *C. elegans*, it is informative to convert the mutant fraction into units of average number of mutant mtDNA molecules per worm (using 959 cells containing a total of ~300,000 mtDNA molecules per worm (Tsang & Lemire 2002a; Gruber *et al.* 2011)). Based on our data, a typical worm on average carries 82 ± 9 (mean ± 1 SD over all 3 age groups) mtDNA molecules with a deletion, irrespective of age. This indicates that mtDNA deletions are extremely rare; less than 0.1 mtDNA deletion per cell on average. Importantly, even under the strong assumption that all of these deletions are concentrated in a single cell, the mutant fraction in that cell would on average still be ~27%, well below the 60% threshold commonly assumed for physiological relevance in mammals (Rossignol *et al.* 2003). Furthermore, in comparison to mammals, C. elegans have been reported to be tolerant towards high levels (up to 80%) of inherited mutant mtDNA with little ill effect and can remain viable even with the homozygous loss of mitochondrial polymerase enzyme (Tsang & Lemire 2002b; Bratic *et al.* 2009). This means that the observed levels of mtDNA deletion would likely have very limited, if any, physiological consequences. These data also raise the question if mtDNA deletions are sporadic (occur independently of each other) or are clonal (arise by clonal expansion from one original mutation and within a single cell).

**Figure 5.**
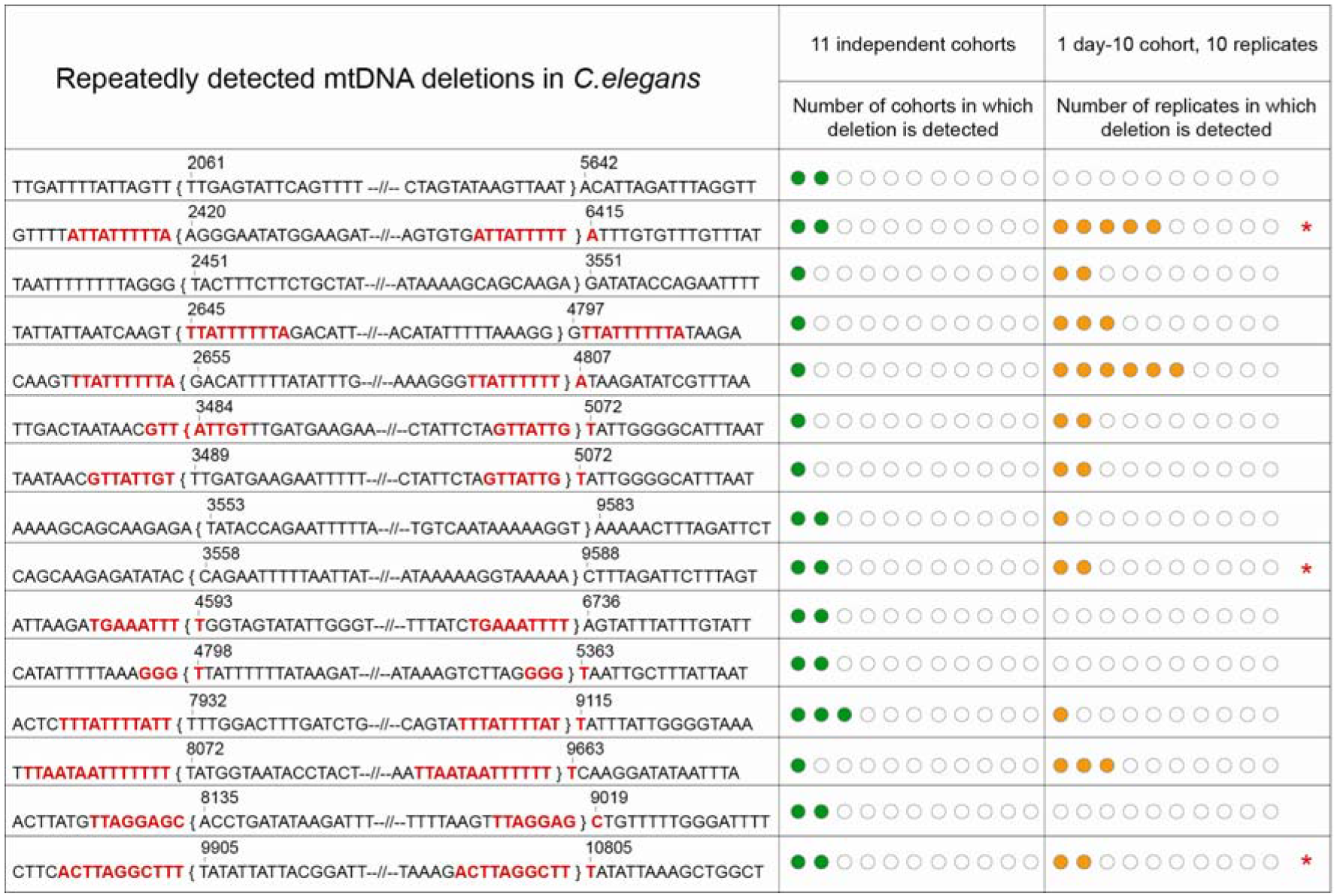
List of all mtDNA deletion sequences that were detected more than once, either within the same cohort or between cohorts. Breakpoint positions, mtDNA sequence near breakpoints and presence of direct repeats (highlighted in red) are shown for each mtDNA deletion that was detected more than once. Sequence positions within the curly brackets are lost by this deletion. Only 15-16 nucleotides near the breakpoints are shown in the figure with the majority of the lost nucleotides in between the breakpoints being replaced by “--//--“. The number of green circles indicates the number of cohorts in which a deletion was detected within 11 independent cohorts. The number of orange circles indicates the number of replicates in which a deletion was detected within the 10 replicate mutation capture experiments of a single day-10 cohort. Deletions marked with red asterisks were detected repeatedly both among cohorts as well as among replicate experiments for the day-10 cohorts.

### Discriminating between clonal expansion and mutation hotspot hypotheses

To examine the clonal nature of mtDNA deletions, we analyzed the deletion spectra detected in 11 nematode cohorts for repetitively detected mtDNA deletions. Clonal expansion would result in substantial increase in copy number of a single deletion originally deriving from a specific deletion event. Such amplification of individual deletions would be supported by repeated observations of the same mutation sequence among replicate samples taken from the same cohort. However, repeated occurrence of deletion breakpoints at a specific mtDNA position can also occur if that region is a mutation hotspot, e.g. this is the case for the 13 bp direct repeat (DR) and 16070 bp position hotspot in human mtDNA (Krishnan *et al.* 2008). Mutation hotspots can arise due to sequence features that promote deletion formation (Nicholas *et al.* 2009). Repeat detections deriving from either a mutation hotspot or from non-hotspot clonal expansion can be discriminated from each other based on the nature of their repeated detection patterns (Nicholas *et al.* 2009). A hotspot deletion is expected to occur independently in different cohorts but may also be repeatedly detected within a single cohort. By contrast, a clonally expanded deletion present in one cohort is unlikely to also be detected in multiple independent cohorts (unless it is also a hotspot).

In total, we sequenced 312 mtDNA deletions, yielding 266 unique deletion sequences detected from eleven independent cohorts (see Table 1). Clonally expanded deletions are typically absent in young tissues and are detected usually in aged tissues (Bua *et al.* 2006). Hence, to maximize the chance of detecting the presence of clonally expanded mtDNA deletions, we performed ten replicate mutation capture experiments using mtDNA extract from one old (day-10) cohort. To evaluate the evidence for clonally expanded mtDNA deletions, we compared the number and breakpoint sequences of deletions that were repeatedly detected within the ten replicate experiments for the single day-10 cohort with those of deletions detected repeatedly among the all 11 independent cohorts (Fig. 5).

Out of the total 266 unique deletions detected, only nine deletions were detected independently in more than one cohort (Fig. 5; also see Table S3, Supporting information). Among these nine deletions, eight were detected in exactly two independent cohorts and one was detected in three independent cohorts. Interestingly, six of these nine deletions have DRs directly flanking their breakpoints, supporting the notion that these deletions may have arisen due to hotspots (Fig. 5). This interpretation is consistent with the fact that these deletions occurred in independent cohorts. Within the single day-10 cohort, we detected a total of 148 deletions, forming a set of 124 unique deletion breakpoint sequences. Nine deletions (i.e. 7.26%) were detected more than once within this day-10 cohort but the vast majority of deletions (i.e. 92.74%) were detected exactly once within the ten replicate experiments. Seven out of the nine deletions that were repeatedly detected within the single day-10 cohort again had long DRs (size ≥ 8bp) exactly flanking their breakpoints, supporting the notion that their repeated occurrence may have been aided by hotspots. This is also the case for all three deletions that were repetitively detected both within the day-10 cohort and between cohorts (Fig. 5). Only two deletions were detected with relatively high frequencies within the day-10 cohort (5 and 6 detections, respectively). Both of these deletions are associated with flanking DRs and one of them was also independently detected in one other independent cohort, again suggesting that mutation hotspots may have contributed. However, we cannot exclude the possibility that some clonal expansion did also occur in these cases (Fig. 5). Nevertheless, overall, the number of repeatedly detected deletions is low both within- and between-cohorts. These data suggest that the majority of the deletions are sporadic and remain at low abundance even in old animals.

### Stochastic model-based analysis of mtDNA deletion accumulation in ageing *C. elegans*

Given that clonal expansion of mtDNA deletions in humans typically takes decades before physiologically relevant levels of expanded deletions can be detected (Bua *et al.* 2006), one possible explanation for the low abundance of mtDNA deletions and the lack of evidence for clonal expansion might be that the lifespan of *C. elegans* is simply too short to allow for efficient amplification of mtDNA deletions. To test this explanation, we employed a stochastic model based on one that we have previously developed for mtDNA deletion dynamics in mice (Poovathingal *et al.* 2009). This computational model is a stochastic model tracking the emergence and evolution (loss or expansion) of mtDNA deletions using realistic dynamical parameter values for mtDNA turnover and mutation rate (Poovathingal *et al.* 2009). We have previously used this model successfully to explore aspects of the dynamics of mtDNA deletions in the *polg* “mutator mouse” and to explore the role of key dynamical parameters for mtDNA deletion clonal expansion (Poovathingal *et al.* 2009; Poovathingal *et al.* 2012). To apply this model to ageing nematodes we scaled relevant parameters to reflect differences between mice and nematodes, including addition of a detailed model of cell division during nematode development. The two key parameters are the rate at which new mutations arise (*de novo* mutation rate) and the rate at which mtDNA molecules are degraded and replicated (i.e. the turnover rate or mtDNA half-life). We considered the possibility that high *de-novo* mutation rate and fast turnover might allow mtDNA deletions to expand to physiologically significant levels even over the short lifespan of *C. elegans. De-novo* point mutation rate is related to fidelity of the DNA polymerases and the extent of oxidative damage in the template DNA (Alexeyev *et al.* 2013). The rate of occurrence of mtDNA deletions *in vivo* is unknown, but the rate of occurrence of the frequently observed human mtDNA ‘common deletion’ has been estimated to be 5.95×10^−8^ per mtDNA replication in cell culture (Shenkar *et al.* 1996). We have conservatively assumed that mutations occur at a much higher rate in *C. elegans* than mammals and have therefore chosen an extremely high *de-novo* mutation rate (up to 2×10^−4^ per mtDNA replication or more than 3000 times higher than the above reference value). This value is extremely conservative in that higher mutation rates promote deletion formation and exaggerate the rate of decline in mtDNA integrity. By choosing a value that is substantially higher than the mutation rates commonly assumed in mice and humans or any suggested mutation rate found in the literature, we therefore establish an upper limit for the amount of impact that mtDNA deletions can have on the age-dependent functional decline in *C. elegans*.

**Figure 6.**
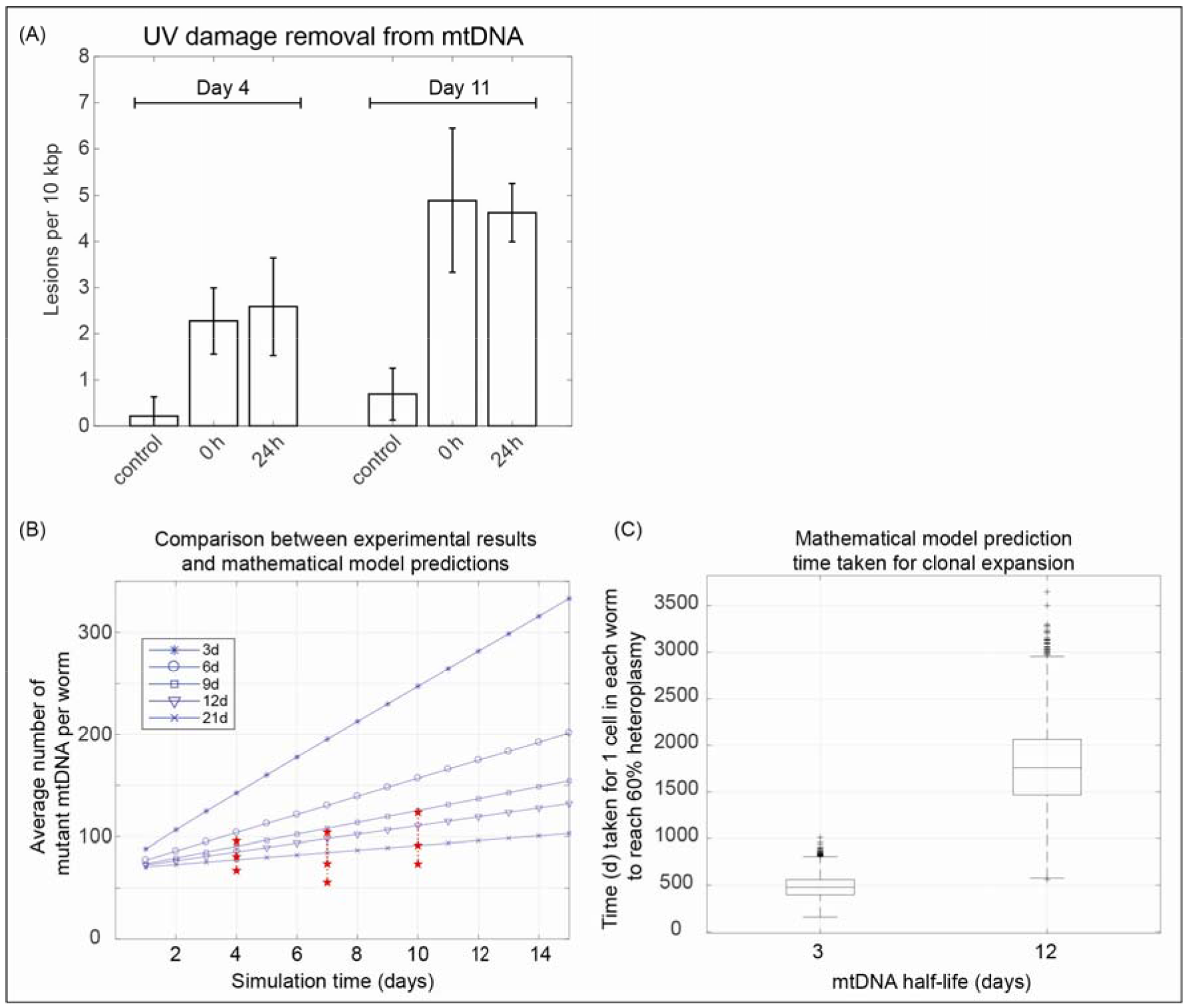
Mitochondrial DNA turnover and clonal expansion of deletion mutations in *C. elegans*. (A) UV induced mtDNA damage levels in control, 0 h and 24 h samples following exposure to 400J/m^2^ of UV radiation at day-4 and day-11 of age. Older animals are more sensitive to UV damage but for both day-4 and day-11 there is a significant increase in mtDNA damage upon exposure to UV (pre-exposure control vs. immediate post exposure). However, UV damage levels are not statistically significantly different between 0 h (immediately post exposure) and 24 h samples (t-test, p-value ≥ 0.18), indicating only a negligible amount of UV damage is removed by mtDNA turnover within 24 h. This rules out mtDNA half-life values of 24h or less and instead suggests half-life values on the order of days (B) Comparison of model predictions for different mtDNA half-life parameter values with experimental data on observed mtDNA mutation burden. Mutant mtDNA molecule counts per worm predicted by the stochastic model (blue) for extremely high mutation rate and various mtDNA half-life values are overlain with the 95% confidence intervals of observed mutant burden based on mutation capture assay (red). Observed mutation burdens are incompatible with half-life values below 9 days. (C) Simulation results estimating the time required until the first cell in an average nematode would reach a physiologically relevant intracellular mtDNA mutant fraction (threshold of 60%) by clonal expansion. This time is highly dependent on mtDNA turnover (half-life), but even for a half-life of 3 days it significantly exceeds the lifespan of *C. elegans*.

### Estimation of mtDNA half-life values in *C. elegans*

There is considerable uncertainty regarding mtDNA half-life values *in vivo* (Poovathingal et al. 2012). One way to establish an upper limit for mtDNA turnover rates is by experimentally tracking the removal of DNA damage from mtDNA. Oxidative damage is removed in mtDNA both through specific DNA damage repair systems as well as by degradation and re-synthesis (turnover) of mtDNA molecules. UV lesions, unlike oxidative damage, are not enzymatically repaired within mtDNA because mitochondria lack nucleotide excision repair required for repairing UV induced damage (Alexeyev *et al.* 2013). Similarly, ethidium bromide (EtBr) intercalation cannot be repaired by the mtDNA repair machinery (Rooney *et al.* 2014). Thus, UV damage and EtBr intercalation can only be removed from mtDNA by degradation and re-synthesis of mtDNA. The dynamics of UV and EtBr lesions thus depend directly on mtDNA turnover rate and their clearance can be used to directly estimate the upper limits of mtDNA turnover (Bess *et al.* 2012; Rooney *et al.* 2014).

We found 2 reports in the literature regarding the rate of UV lesion removal from mtDNA in *C. elegans* (Meyer et al. 2007; Bess et al. 2012) and we also carried out our own experiments, using our sequence specific mtDNA damage assay to observe the repair kinetics of UV induced lesions (Fig. 6A). Based on the first order rate of decline in mtDNA following EtBr treatment, mtDNA half-life values of between 8 and 13 days have been reported for *C. elegans* (Rooney *et al.* 2014). The same study also reports a mtDNA half-life value of 5 days based on the kinetics of UV damage removal (Bess *et al.* 2012; Rooney *et al.* 2014). Assuming a first order decay kinetic we re-analyzed data on *C. elegans* mtDNA UV damage removal from Meyer et al. (Meyer *et al.* 2007) and obtained a shorter half-life value of 3 days for mtDNA. In our own experiments, we found that there was no significant repair of UV induced damage after 24 hours (Fig. 6A). In fact, we could not find a significant decrease in UV damage at 24 h when compared to immediately after UV exposure, indicating that the turnover/removal of UV damaged mtDNA is too slow to result in a significant loss of UV lesions within 24h. This, again, suggests a mtDNA half-life of at least several days. Based on these data we set a range for mtDNA turnover of between 3 days and 12 days. We then simulated mtDNA deletion dynamics using mtDNA half-life values between 3 days (fastest rate as suggested by the above estimates) and 12 days (best estimate) together with a conservative (very high) level of mtDNA mutation rate of 2×10^−4^ per mtDNA replication. We find that our stochastic model agrees best with our experimentally observed mutation burden in ageing worms for a half-life value of 9-21 days (i.e. slow turnover) (Fig. 6B). Using shorter half-life values, we find that the model prediction is qualitatively incompatible with the observed mutation burden (Fig. 6B), suggesting that half-life values for *C. elegans* mtDNA are unlikely to be 3 days or lower if the mutation rate is at 2×10^−4^ per mtDNA replication.

### Estimating the time required for clonal expansion in *C. elegans*

However, even assuming an extremely fast turnover rate (a half-life of 3 days) and a very high mutation rate (2×10^−4^ per mtDNA replication), the predicted mtDNA deletion burden still remained well below the ~60% threshold required for ETC inhibition throughout the life of *C. elegans* (Fig. 6B). Most importantly, detailed examination of the simulation trajectories revealed that, while overall mutation burden increases with a faster turnover, there are essentially no cells with clonally expanded mtDNA, even for parameter values most favorable for clonal expansion (very fast turnover and very high mutation rate). To illustrate this lack of clonal expansion we used our model to evaluate how long it would take in an average nematode worm for the first cell with high abundance of clonally expanded mtDNA to emerge. Using a half-life value of 12 days, the value most consistent with UV and EtBr lesion dynamics, it would take on average about 4.8 years for the first cell in the average nematode to cross this critical threshold of clonal expansion (Fig. 6C). Even with a more rapid, physiologically less plausible half-life value of 3 days, it would take 1.37 years for the first cell in the body of a *C. elegans* worm to reach the threshold of biochemical compromise (Fig. 6C). This may explain why mtDNA deletions remain rare even in old nematodes and why there is no strong evidence for clonal expansion from our experiments. The data suggest that, under the assumptions of our model, nematode lifespan is at least an order of magnitude too short for clonal expansion of mtDNA deletions to play any role in nematode ageing.

## Discussion

Mitochondrial decline and dysfunction are common features of ageing and of many age-dependent diseases in mammals and in model organisms including nematodes (Braeckman *et al.* 2002; Herndon *et al.* 2002; Yasuda *et al.* 2006; Gruber *et al.* 2011; López-Otín *et al.* 2013). In humans, one of the mechanisms that is believed to be involved in mitochondrial ageing is loss of mtDNA integrity and accumulation of mutant mtDNA (Stewart & Chinnery 2015). The accumulation and clonal expansion of mutant mtDNA are processes that have been causally linked to neurodegeneration and sarcopenia (Bender *et al.* 2006; Herbst *et al.* 2016). Metabolic decline, mitochondrial dysfunction and sarcopenia are all conserved features of ageing in *C. elegans* (Braeckman et al. 2002; Herndon et al. 2002; Fisher 2004; Yasuda et al. 2006; Gruber et al. 2011). While some evidence for an age-dependent increase in mtDNA deletion probability has been reported in *C. elegans* over 20 years ago, the question if these deletions are functionally linked to metabolic and mitochondrial ageing has remained unanswered (Melov *et al.* 1994; Melov *et al.* 1995). Using cohorts of ageing nematodes and 236 distinct primer pairs, we comprehensively determined the mtDNA deletion spectrum and mutation burden in ageing *C. elegans*, identifying 266 novel mtDNA deletions (see Supporting document 2). Partially duplicated mtDNA (pd-mtDNA), which are large circular mtDNA molecules containing one deleted molecule and one wild type molecule, have been reported in mitochondria (Holt *et al.* 1997). Differentiating a pd-mtDNA from a mtDNA deletion is challenging and can require a large DNA sample, single cell data and specific PCR design (Bodyak *et al.* 2001). Our mutation capture method is not designed to identify pd-mtDNA. The abundance of any specific type of pd-mtDNA is suggested to be low (Bodyak *et al.* 2001) and hence, for simplicity, we assume that all the identified breakpoint junctions are deletions and not duplications. However, presence of pd-mtDNA within our dataset, if any, would only reduce the mtDNA deletion fraction further and would therefore not affect the key conclusions from this study. On average, we detected a slightly larger number of deletions per experimental replicate in aged nematode cohorts compared to young cohorts (Table 1), a finding that is consistent with the only previous report (Melov *et al.* 1995). For quantification, we employed Monte Carlo simulations, taking into account the experimental design and sampling statistics. We find that the apparent age-dependent increase in mutation burden is not statistically significant and that overall burden remains low over the course of life, with on average 82 ± 9 (mean ± 1 SD over all 3 age groups) mutant mtDNA molecules per worm irrespective of age. This mutation burden is well below the 60% threshold where mtDNA deletions would become physiologically limiting in terms of mitochondrial energy production (Rossignol *et al.* 2003). Even under the extremely unlikely assumption that all mutant molecules in each worm were concentrated in a single cell, the levels of mtDNA deletion in even that single cell remain under this 60% threshold. Under the much more plausible assumption that these ~80 mutations are relatively equally distributed amongst the 959 cells of an adult nematode, these data suggest that most cells in *C. elegans* harbors no mutant mtDNA and that, therefore, mtDNA mutations at this level are unlikely to be sufficient to cause significant physiological detriments. We determined deletion burden up to day 10 of life because at a temperature of 25°C, this is an age where rapidly rising mortality, functional and metabolic decline and loss of mitochondrial integrity suggest significant age-dependent failure (Gruber *et al.* 2011).

Interestingly, worms with high (20-80%) mutant burden of inherited *uaDf5* mtDNA deletion have been reported to be free of major defects (Tsang & Lemire 2002b) although they have been reported to exhibit reduced fitness and sperm performance (Liau *et al.* 2007). Flies have also been shown to tolerate high burden of mtDNA deletions without serious functional compromise (Petit *et al.* 1998; Tsang & Lemire 2002b). Based on these findings, it has been suggested that simple organisms such as worms and flies can tolerate high burden of mtDNA deletions, further supporting the notion that the low levels detected by us are unlikely to be detrimental (Tsang & Lemire 2002b). This suggests that mtDNA deletions at the level that we detect are unlikely to play any functional role in nematode ageing.

Furthermore, if clonal expansion was a prominent process during the ageing process of *C. elegans*, we would expect higher abundance, and hence repeated detections, of clonally expanded deletions. This should have been detected during within-cohort replicate experiments relative to between-cohort comparisons. However, number of repeatedly detected mtDNA deletions was low in both within- and between-cohorts. Furthermore, majority of these repeatedly detected deletions were associated with flanking direct repeats in the mtDNA sequence, a feature that has been frequently associated with mtDNA deletions (Lakshmanan *et al.* 2012; Lakshmanan *et al.* 2015).

Our stochastic model provides a possible explanation for the relatively low abundance of mtDNA deletions and absence of evidence for clonally expanded mtDNA deletions in ageing nematodes. The computational model shows that, even using extremely conservative assumption (very high mutation rates and very fast mtDNA turnover rates), nematode lifespan is likely too short to allow for clonal expansion to occur. This finding may not be surprising given that clonal expansion in human typically takes decades before significant mutation burden is detectable, and even then only in a small number of cells (Elson *et al.* 2001; Kowald & Kirkwood 2013). To compress dynamic trajectories spanning half a century in humans into less than a month in nematodes would require extreme and biologically unrealistic parameter values to force clonal expansion to proceed roughly 600 times faster in *C. elegans*. Moreover, according to one study, even in muscle of a 92-year-old individual less than 1% of skeletal muscle fibers exhibited electron transport chain abnormalities and clonally expanded mtDNA deletions (Bua *et al.* 2006). For clonal expansion to significantly impact the 302 neurons or 187 muscle cells of an average adult *C. elegans*, let alone to explain the general decline in metabolic rate seen in ageing nematodes (Braeckman *et al.* 2002; Yasuda *et al.* 2006; Gruber *et al.* 2011), the fraction of affected cells would have to far exceed the level seen even in the oldest humans (up to 30% of neurons) (Itoh *et al.* 1996; Kraytsberg *et al.* 2006). For clonal expansion to impact a significant fraction of nematode cells, we would have to assume mtDNA half-life values far shorter than even the most rapid mtDNA turnover rate previously suggested for any animal. Such extreme values are also inconsistent with our and the publically available data on the removal rate of oxidative, UV and EtBr lesions in nematode mtDNA (Fig. 6).

In summary, we find experimentally that *in C. elegans* (1) there is no statistically significant increase in mtDNA deletion burden with age and (2) the mutant burden, even in old animals, is less than would be required to significantly affect cell respiration. The levels of deletions observed even in aged nematodes appear too low to be physiologically relevant and certainly are too low to explain the robust age-dependent decline in metabolic rate seen in ageing *C. elegans*. Furthermore, there is no strong evidence that clonal expansion of mtDNA deletions commonly occurs in *C. elegans* and our stochastic modelling results suggest that nematode lifespan is simply too short for clonal expansion to occur in most ageing nematodes. Together, our findings suggest that mtDNA deletions do not play a significant role in age-dependent functional and metabolic decline in nematodes and that mtDNA deletion dynamic does not determine nematode lifespan.

Previous experimental confirmation of clonal expansion of mtDNA deletions in muscle and brain has mostly been in long-lived animals, typically humans, monkeys and less frequently in rat. By contrast, while some deletions have also been reported in aged mice, several experimental studies have indicated a paucity of mtDNA deletions even in aged mice tissues (Williams *et al.* 2010; Ameur *et al.* 2011). These and our data are consistent with two previous mathematical models of mtDNA deletion dynamics that also concluded that such models did not produce clonally expanded mtDNA deletions within the lifespan of mice (Kowald & Kirkwood 2013; Kowald *et al.* 2014).

It is therefore possible that mtDNA deletion accumulation and clonal expansion is not a public mechanism of ageing but one that is private to relatively long-lived animals. However, it is important to remember that, by contrast, structural and functional deterioration of mitochondria and general hypo-metabolism all are key public features of ageing that are well replicated in nematodes. Our findings suggest that the causes of these conserved features of mitochondrial ageing may not be mtDNA deletion accumulation but other, possibly public, mechanisms of ageing.

## Experimental Procedures

### C. elegans strain and maintenance

JK1107 (*glp-1*) was cultivated and maintained at 25 °C to prevent progeny. Preparation of nematode growth media was as previously described (Stiernagle 2006) with the addition of Streptomycin (Sigma) at the final concentration of 200 μg/ml. Streptomycin-resistant bacteria strain (Escherichia coli OP50-1) was added to the solid media at 10^10^ cells/ml.

### Extraction and purification of mtDNA

Adult worms were harvested on days 4 (young), 7 (middle-aged) and 10 (old). Worms were washed off plates using ice-cold M9 buffer (Stiernagle 2006) and allowed to settle on ice, washed twice with ice-cold S-basal buffer to remove bacteria, and twice with ice-cold isolation buffer (210 mM mannitol, 70 mM sucrose, 0.1 mM EDTA, 5 mM Tris-HCl, pH 7.4). Worms were homogenised in isolation buffer on ice, using a Teflon pestle homogeniser. Homogenate was sampled and checked under light microscope to ensure complete homogenisation. Worm homogenate was centrifuged at 600 g, 4 °C for 10 min to remove cell debris. Supernatant was centrifuged at 7,200 g, 4 °C for 10 min to obtain mitochondria. Pellet was re-suspended in TE buffer (50 mM Tris-HCl, 0.1 mM EDTA, pH 7.4). Mitochondrial DNA was treated with PrepMan Ultra Sample Preparation Reagent (Thermofisher Scientific) according to the manufacturer’s protocol.

### Mitochondrial DNA copy number quantification

Mitochondrial copy number was quantified according to (Schaffer *et al.* 2011). Briefly, individual worms were picked into PCR tubes, lysed, and mtDNA copy number of individual nematodes was determined by quantitative real-time PCR (qRT-PCR) as previously described (quantified by serial dilution) (Gruber *et al.* 2011; Schaffer *et al.* 2011).

### Random deletion detection qPCR assay

Random deletion detection qPCR assay was performed using KAPA PROBE FAST qPCR Kits (KAPA Biosystems) on ABI StepOnePlus and ViiA 7 real-time PCR systems (Life Technologies). Each real-time PCR reaction contained: 8.19 μl nuclease-free water (Promega), 10 μl KAPA PROBE FAST qPCR Master Mix (1X), 0.18 μl Forward Primer (900 nM, refer to list of primers in Table S4, Supporting information), 0.18 μl Reverse Primer (900 nM), 0.05 μl Taqman Probe (250 nM), 0.4 μl ROX Reference Dye (500 nM), 1 μl sample (mtDNA or water). The real-time PCR cycling protocol was enzyme activation at 95 °C for 3 min, then 60 cycles of denaturation at 95 °C for 3 sec and annealing/extension/acquisition at 60 °C for 1 min 15 sec. Mutation capture method preferentially amplifies the mutant mtDNA molecules carrying deletion (Fig. S5, Supporting information). Each mutation capture experiment involved 236 distinct PCR reactions (1 PCR per primer pair). Equal volumes of diluted mtDNA stock solution was aliquoted into 236 wells based on preliminary experiments. Successful amplification indicates the likely presence of mutant and samples from fluorescent wells were sequenced to confirm and identify the breakpoint positions of the mtDNA deletion. A set of positive and negative controls were used during each mutation capture assay. Detailed description of the mutation capture assay plate design, controls used and the results for controls can be found in the Supporting Information (see Fig. S6).

### Validation and sequencing of PCR amplification product

Each putative deletion (amplified well) was subjected to agarose gel electrophoresis to check for presence and size of a deletion fragment. 15 μl of PCR reaction from each well was then purified using QIAquick PCR Purification Kit (QIAGEN) or, in cases with more than one product, gel DNA extraction using QIAquick Gel Purification Kit (Qiagen). Products were sequenced and breakpoints confirmed using Nucleotide BLAST by aligning sequences of the fragments to the *C. elegans* wild type mtDNA sequence (www.ncbi.nlm.nih.gov/blast/), NCBI accession number: X54252.1/WBcel235.

### Induction of UV damage and repair assay in mtDNA in live nematodes

Synchronous cultures of day 4 and day 11 worms (about 10,000 worms per condition) were washed off the plates. The worms were washed three times with M9 buffer to remove excess bacteria. Worm pellets without bacteria were transferred to fresh NGM agar plates without food. The worms were evenly spread across the plates to prevent overlapping of worms. The worms were irradiated using a CL-1000 ultraviolet crosslinker with an emission peak at 254 nm. Worms on NGM agar plates without the plates cover were exposed to 400 J/m^2^ UVC source. Immediately after UVC exposure, worms exposed to UVC without any repair were washed off the plates and collected for the mtDNA extraction and purification. Another 10000 of UVC irradiated worms were transferred to fresh NGM agar plates seeded with OP50-1, the worms were allowed to grow and repair any damage for 24 hours. Worms were harvested for mtDNA extraction after 24 hours.

### mtDNA damage assay

PCR based damage quantification was done as described elsewhere (Melov *et al.* 1995; Gruber *et al.* 2011; Qabazard *et al.* 2014). Briefly, following mtDNA extraction and purification, mtDNA damage was assessed using two independent qRT-PCRs: short-qRT-PCR and extra-long qRT-PCR. The amplification efficiency in 6.3 kb region (forward primer: 5’-TCGCTTTTATTACTCTATATGAGCG-3’ and reverse primer: 5’-TCAGTTACCAAAACCACCGATT-3’) of mitochondrial genome (Melov *et al.* 1995) was compared to a 71 bp region (forward primer: 5’-GAGCGTCATTTATTGGGAAGAAGA-3’ and reverse primer: 5’-TGTGCTAATCCCATAAATGTAACCTT-3’). Short qRT-PCR was performed using FAM-labeled Taqman probe and TaqMan Universal PCR Master Mix according to manufacturer’s instructions (Applied Biosystems); extra-long qRT-PCR was conducted using SYBR Green I dye and GeneAMP XL PCR kit according to manufacturer’s instructions (applied Biosystems). Briefly, Short-qRT-PCR was conducted under the following cycling conditions: 2 min at 50 °C, 10 min at 95 °C, 50 cycles of 15 s at 95 °C and 60 s at 60°C. Extra-long qRT-PCR was performed on the same Real-Time PCR system using the following cycling conditions: 2 min at 94 °C, followed by 60 cycles of 10 s at 93 °C, 10 min at 62 °C, and 30 s at 68 °C, ending with a final extension of 30 min at 68 °C. mtDNA lesions for the samples were then quantified from the ct values as previously described (Neher & Sturzenbaum 2006; Santos *et al.* 2006; Meyer 2010; Gruber *et al.* 2011; Ng *et al.* 2014).

### Stochastic model of mtDNA mutation dynamics

The stochastic modelling framework used in this work to study mtDNA mutation dynamics is identical to the mathematical model developed previously by our group (Poovathingal *et al.* 2009). *C. elegans* specific developmental cell divisions and mtDNA copy numbers were used to adopt this previously published model to worms. Complete set of model parameters used for the simulations could be found in the supporting information (see Table S5). Detailed description of model equations and parameters could be found in our original model publication (Poovathingal *et al.* 2009).

## Acknowledgments

This work was funded by the Ministry of Education Singapore (Grant R-184-000-230-112) and (Grant MOE2014-T2-2-120).

## Author Contributions

LNL, ZL and LF devised, designed, and performed experiments and analyzed results. JG, RG and BH participated in the experimental design and in the interpretation, conceived the project and coordinated its execution. All authors commented and approved the manuscript.

## Supporting Information List

- Figure S1. Coverage by primer pairs
- Figure S2. Artificially synthesized mutant sequence
- Figure S3. Serial dilution experiments
- Figure S4. Estimation of mtDNA deletion mutation fraction
- Figure S5. Distribution of mutant mtDNA amplicon lengths
- Figure S6. Plate design
- Table S1. Experimental design
- Table S2. Results from replicate mutation capture experiments
- Table S3. Repeatedly detected mtDNA deletions
- Table S4. Primer positions and sequence information
- Table S5. Parameter values used for stochastic model simulations

